# A Pumpless, High-Throughput Microphysiological System Confirms Enteric Innervation of Duodenal Epithelium Strengthens the Barrier Function

**DOI:** 10.1101/2023.06.03.543561

**Authors:** Kyla N. Nichols, Jessica R. Snyder, Ryan A. Koppes, Abigail N. Koppes

## Abstract

Enteric neurons, diverse in function and great in number, are heavily involved in homeostasis within the small intestine and their dysregulation has been implicated in gastrointestinal disorders and neurodegenerative diseases. Innovations in biofabrication have resulted in advances for in vitro models of the gut, however the majority lack enteric innervation, limiting therapeutic screening and discovery. Here, we present a high-throughput co-cultured microphysiological system (MPS), or organ chip, that supports a primary epithelial monolayer that directly interfaces with a three-dimensional hydrogel containing a primary enteric neuron culture, mimicking the close proximity present in vivo. The acrylic MPS device was fabricated with our established and cost-effective laser cut and assemble method. We have expanded this technology to include up to twelve 3D MPSs per device within the footprint of a traditional well-plate, supporting high-throughput experimentation. The inclusion of this 3D microtissue does not hinder physiologically relevant flow, standard measures of barrier function, and microscopy techniques. The device features gravity-driven flow to induce physiological shear stress on the epithelium culture and provide continuous nutrient presentation. Results show the intestinal and neural tissue maintained expected morphologies over an experimental timeline of ten days. Proximal enteric neurons extend neurites through the 3D hydrogel towards the epithelial monolayer. Barrier function was confirmed with both Transepithelial Electrical Resistance (TEER) and Lucifer Yellow diffusion on-chip. TEER confirmed a significantly more substantial barrier integrity in co-cultures compared to baseline values (1.25-fold) in epithelial cell-only. Lucifer yellow permeability assays performed in parallel supported the TEER results, with an 11.8% lower permeability of the co-cultured group than the epithelium only. The presence of the ENS on chip results in a significant (1.4 fold) reduction in epidermal growth factor (EGF). This is the first high-throughput, innervated gut on a chip device that demonstrates the importance of the autonomic nervous system on EGF expression and possibly epithelial renewal in vitro. Innervation is essential to create more biomimetic and physiologically relevant in vitro models for biological and pharmacological assays.

---

The gastrointestinal (GI) tract forms a selectively permeable barrier, allowing nutrient transport into the host while keeping pathogens out. The gut is host to millions of harmful and symbiotic microorganisms that the intestinal epithelium must keep separate from the inner circulatory system. The integrity of this epithelial barrier is essential for systemic health. A compromised intestinal barrier is implicated in several GI disorders due to bacteria and toxins infiltrating the tissue and causing inflammation. These disorders include the discernible Irritable Bowel Syndrome (IBS) and Inflammatory Bowel Disease (IBD), and are also co-morbid with several diseases including depression and hypertension, suggesting gut health impacts the entire body^1-4^. The underlying mechanisms of GI disorders remain elusive due to the complexity of entire organisms, especially considering the interplay between cognitive and gut health during in vivo investigations. Therefore, a reductionist approach utilizing in vitro models are necessary to understand the regulation of barrier function and the impact the gut can have on systemic health.

Gut microphysiological systems (MPSs), or organ chips, have been developed with various materials and architectures to add physiological relevance, such as human tissue, inflammatory compounds, shear stress to model peristalsis, compositions of the microbiome, and villi-like structures^5-9^. Due to their advantages over in vivo models, these MPSs are used to study broad areas, from drug delivery to disease progression^10,11^. The expanded complexity of gut-on-a-chip systems, including physiological components such as shear stresses via medium perfusion, have shown faster differentiation and increased mucus production compared to traditional 2D culture methods^8,12^. These MPSs also support analysis in real-time, including standard light microscopy and the ability to assess barrier function and permeability, which are especially relevant in modeling epithelial dysfunction such as in IBD. Transepithelial electrical resistance (TEER) and apparent permeability measurements of fluorescent molecule diffusion are non-destructive methods often used to assess barrier integrity of live epithelial monolayers on chip^8,13,14^. The health of these epithelial barrier properties are pivotal for studying microbiome interactions and drug absorption^15,16^. These techniques are easily implemented in transwell cultures, however the difficulty and cost of integrating especially TEER is directly correlated to the complexity of these MPSs. Here, we present a new low-cost platform that breaks the coupling of cost between increased biorelevance and real-time outputs.

Current gut epithelium cultured in these MPSs require cell lines, isolated primary cells, or induced stem cells. The human colorectal adenocarcinoma cancer cell line, Caco-2, is frequently used as an epithelium model with maturation leading to tight junction formation. However, Caco-2 cells have limitations since they only comprise of the absorptive enterocyte phenotype, lacking the heterogeneous population found in vivo^17^. The field is moving to primary epithelium cell models, often seeding monolayers of epithelial cells isolated from expandable intestinal epithelial organoids. Primary organoids proliferate in three dimensions and contain intestinal stem cells and additional epithelial cell subtypes important in neural, immune, and microbial cell interactions including absorptive enterocytes, mucus-producing goblet, enteroendocrine, and Paneth cells^18^. Primary epithelium models are obtainable through dissected rodent tissue or human biopsy samples and proliferate easily, allowing sizable stocks of the cells to be cultured or cryopreserved for up to several years^19^. Degree and synaptic organization of the enteric nervous system (ENS) on the function of each of these epithelial populations is not fully understood. These heterogeneous epithelium populations more closely resemble the human gut than traditional immortalized cancer cell lines and are needed to explore the role of the autonomic nervous system (ANS) of barrier function.

Sensations and coordination of gastrointestinal (GI) activities are facilitated by the underlying neurons of the gut, the ENS as well as branches of the ANS comprised of plexi, ganglia, spinal cord, and cranial nerves. Greater in total neuron numbers than the spinal cord, enteric neurons inform the epithelium’s proliferation through epidermal growth factor (EGF) signaling to enterocytes, the most abundant epithelial subtype^20^. The enteric nervous system also aids in preventing microbial infection by promoting goblet cell mucus production, resulting in a stronger barrier ^21^ . Epithelial – enteric neuron communication has been found to occur through soluble neuronal mediators as well as direct synapses with enteroendocrine cells^22^. One example of paracrine signaling occurs through Vasoactive Intestinal Peptide (VIP) released by VIPergic enteric neurons. VIP promotes proliferation of the epithelial cells, improves the barrier, and increases secretion^23,24^. Conversely, acetylcholine (Ach) released by cholinergic enteric neurons permeability and decrease epithelial proliferation, suggesting that an imbalance between these two neurotransmitters and barrier dysfunction^25,26^. To further recapitulate native tissue, we incorporated primary enteric neurons encapsulated in 3D into a MPS of primary epithelium monolayers.

In addition to incorporating enteric neurons within our model system, we developed a new MPS that addresses a number of the current limitations that plague many gut-chip designs, including low sample throughput, high media consumption, and reliance on syringe pumps to induce shear. Our MPS design is the first to support primary epithelial monolayers, 3D enteric encapsulation, and up to twelve independent samples all within a standard well plate footprint. Uniform pulsatile flow/shear across each sample is accomplished with media reservoirs that support pumpless, gravity-driven flow to induce physiologically relevant shear stresses when placed on a standard laboratory rocker at 2 degrees and 10 rpm. The device is assembled with a glass base that allows for high resolution microscopy through the height of the MPS so that both the living epithelial and neuron populations can be monitored during the experimental time course and analyzed for end-point characterization.

We hypothesized the presence of enteric neurons would alter the epithelial differentiation and maturation timeframe compared to epithelial-only cultures, mimicking function in living beings. Using our custom MPS system, we studied the impact of enteric neurons on epithelial barrier stability with permeability assays, including TEER and lucifer yellow diffusion, and end point immunocytochemistry. Our results indicate for the first time an engineered and high-throughput innervated gut-chip, and that enteric neurons positively influenced epithelial barrier strength and altered growth factor signaling, highlighting the importance of incorporating innervation when developing biomimetic MPS devices.

## Results

### Co-cultures of epithelial cells and enteric neurons are supported by a cost effective, pumpless, high throughput MPS

We developed a new innervated microphysiological system (MPS) of primary duodenal epithelium to investigate the role of enteric neurons on primary epithelial cell permeability (**Fig 1a and 1b**). The 12-unit MPSs are fabricated using laser-cut thermoplastic layers (**Fig. 1c**). PMMA and PET layers facilitate an oxygen impermeable environment and the 12-sample chip can be produced in under a couple of hours for $21 dollars, or less than $2 per unit/sample (**Fig. 1b&c**). The scaled-up device supports 12 co-culture chambers on a single 76 mm by 101 mm chip which fits well within a standard well plate footprint with dimensions around 127 mm by 85 mm. The MPS features a top chamber on a permeable membrane allowing epithelial monolayer adhesion and polarization shown as a cross section (**Fig. 1b**). There is a chamber supporting three-dimensional culture below the membrane where enteric neurons were seeded within a Matrigel® solution (**Fig. 1c**). Figure 1c shows an exploded view of a single MPS and its culture setup.

**Figure 1.**
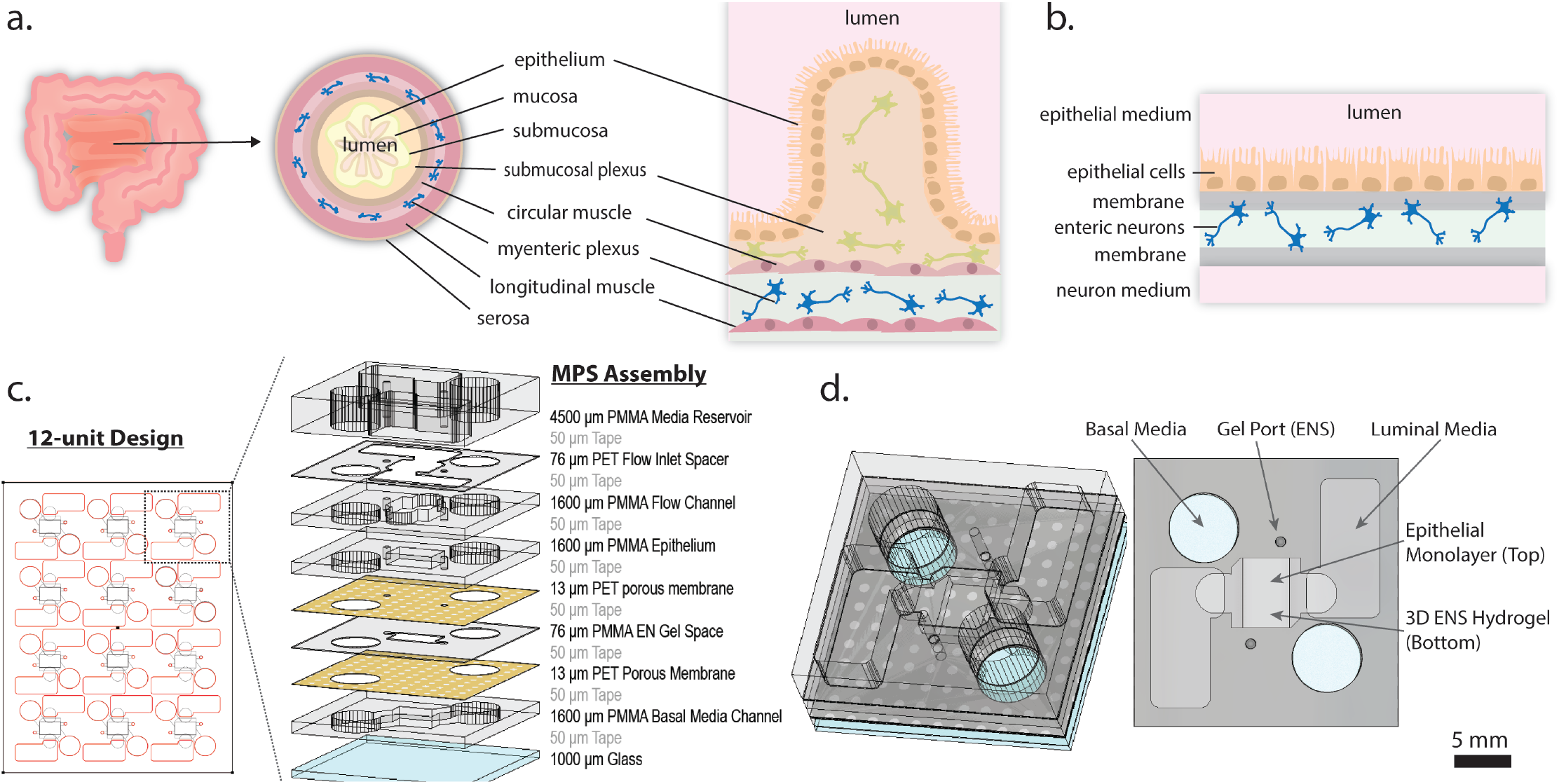
Chip designed to recapitulate the 3D small intestine into a simplified culture system. **a)** Cross sections of the bulk small intestine, microscopic small intestine, and **b)** our reductionist model of the innervated small intestine. **c)** The 12-unit design fits within the dimensions of a standard 12 plate while keeping each 3D MPS independent for up to four experimental parallelization in triplicate. The individual MPS is comprised of laser-cut thermoplastics (PET track etched membrane and PMMA spacers) bonded layer-by-layer with a 3M adhesive tape onto a glass slide. An exploded view of the chip layers and thicknesses. **d)** An orthogonal and straight on assembled version is displayed to highlight cell seeding ports as well as media wells for both the apical and basal chambers.

Optimization of the experimental timeframe was completed to support enteric neuron maturation and extension in the 3D environment of the MPS, as well as epithelial monolayer differentiation (**Fig. 2a**). Neurons were allowed to mature for 7 days before epithelial seeding to allow adequate time for neurite extension throughout the gel layer of the MPS and followed by 3 days in co-culture with the epithelium before endpoint analysis. Twenty-four hours after epithelial seeding, the MPSs were moved to a laboratory rocker to supply shear across the epithelial monolayer, a complete timeline of the experimental design is in Fig 2a.

**Figure 2.**
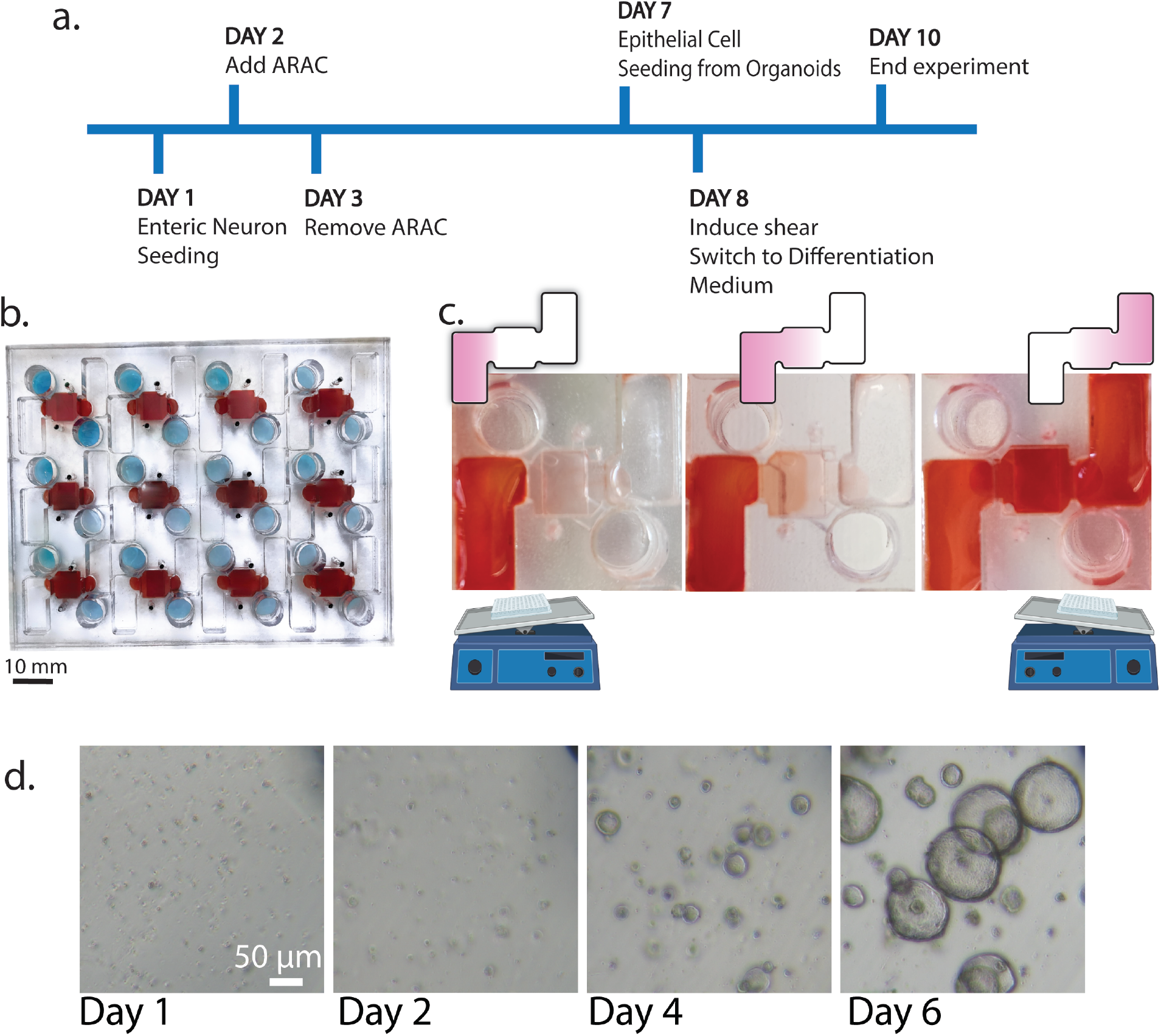
High-throughput chip design allows application of relevant shear stress without the need for tubing or pumps. **a)** Seeding timeline of the MPS with primary rat cells and the duration of experiments were run over 10 days, including 3 days of co-culture. Improvements to our model included **b)** scaling up the device to contain 12 independent culture systems and **c)** supplying gravity-driven flow across the epithelial cells. **d)** Epithelial crypt cells were isolated from the duodenum and expanded as intestinal organoids in a 3D culture environment before use in the experiments as shown in brightfield.

To improve the robustness and ease of the MPS, the model was designed with two large, opposing media chambers to allow the application of relevant shear stress in a pump-free design (**Fig 2**). Gravity-driven flow of medium across the epithelial monolayer was accomplished with a standard platform rocker (**Fig. 2c**). The large plate design eases alignment with the axis of rotation and ensures application of flow across the 12 MPSs. The flowing liquid creates a more biomimetic environment for the intestinal cells than a static culture and avoids the use of pumps and tubing.

Tilt angle and speed were adjusted to achieve physiological, pulsatile shear stress in the range of 0.002– 0.08 dyne/cm^2 6^. Our max shear stress was calculated to

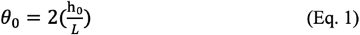

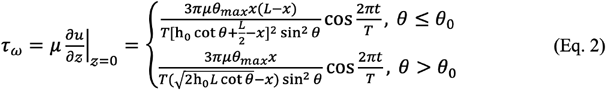

be 0.0683 dyne/cm^2^ across the middle of the cell culture chamber when the rocker was at a maximum tilt angle of 2 degrees and rocking at 10 rpm. Equation 1 was used to calculate the critical angle when the fluid-free surface comes in contact with the edge of the MPS bottom (16.7 degrees). Equation 2 was then used to calculate the max shear stress. Viscosity is a critical component of shear stress. Here, we used a published value of DMEM’s viscosity, 0.731 mPa*s^32^. The shear calculation used several assumptions, including a no-slip boundary condition at the bottom of the dish, a zero-velocity gradient at the fluid-free surface, flow mainly from gravity, and the effect from gravity is much greater than viscous effects. In addition, centrifugal force was neglected due to slow angular speed, velocity was assumed to be normal to the dish bottom, and the pressure gradient along the fluid depth was ignored^28^.

Primary neonatal rat enteric neurons were cultured in three-dimensions in the MPS. Epithelial monolayers were seeded from rat neonatal duodenal organoids (**Fig. 2d**). Using neonatal intestines allowed us to look at early epithelial development and differentiation. Most organoids maintained a spheroid morphology with limited budding up to seven days before passage. Seven days post neuron seeding, epithelial cells from organoids were seeded into the top chip layer chip to form a monolayer. Epithelial cells received a differentiation medium 24 hours after their seeding in the chip. Following the tenth day from initial neuron seeding, all experiments were ended. At the endpoint, the monolayers were no longer completely confluent. Mature epithelial cells have a short lifespan and begin to lift off the chip, similar to anoikis in the body where they shed from the intestinal lining after 2-6 days,^33^ limiting the time frame of these experiments without a source for intestinal renewal.

### Epithelial cell adhesion and enteric neurite extensions occur in an interfacing co-culture MPS

The high throughput chip design was utilized for all cell culture experiments. Healthy cell morphologies; adherent epithelium and spindly neurons, were confirmed using an inverted light microscope prior to running permeability assays and immunostaining. The glass bottom and transparent membranes within the chip allowed for clear imaging through the different layers of the co-culture. Figure 3a and b show the brightfield images of the epithelial monolayer on the top chamber and a plane of the gel containing the enteric neurons before fixation. The epithelial cells developed a cobblestone morphology adhering to the ECM-coated membrane, unlike the budding spheres they form while expanding. The enteric neurons developed long and distinct neurite extensions, forming ganglionic-like morphology confirmed with immunocytochemistry (**Fig. 3b&d**). Mature epithelial cells were demonstrated by expression of ZO1 and a characteristic cobblestone pattern bordering the cells (**Fig. 3c&d**). However, the cell surface did not maintain a uniform matured epithelium, exhibited by some areas of cytoplasmic ZO1 staining. This is likely due to enterocytes’ fast cell turnover period within the duodenum and three-day timeframe for experiments. Figure 3d also features beta III tubulin staining within the neuronal compartment of the MPS, highlighting the presence of mature enteric neurons in the co-culture.

**Figure 3.**
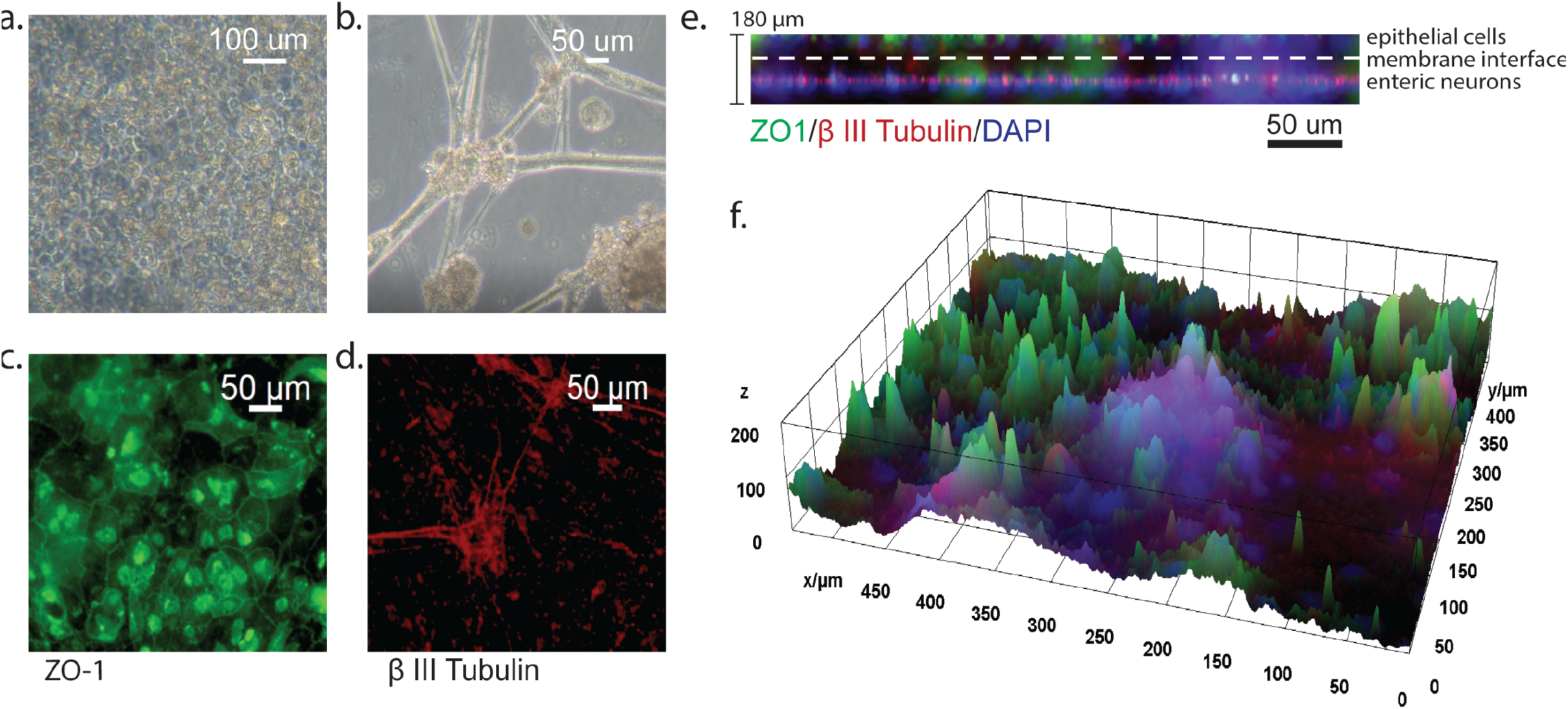
Characterization of epithelial barrier and ENS interaction on-chip. **a)** Representative brightfield images (day 10) of living epithelial and **b)** neuronal layers of the organ chip before fixing and staining. **c)** Immunostained layers of the MPS feature the epithelium, showing the ZO-1 tight junctions in the characteristic cobblestone pattern, and **d)** mature neuron cultures stain with beta III tubulin. These images were taken in the same location at different focal planes in the z-direction. **e)** An orthogonal maximum intensity projection shows the layers of the culture, beta III tubulin identifying the neurons and ZO-1 the epithelium. (f) A 3D surface plot showing the proximity of the different cell types in the culture systems and the depth.

A unique feature of this system is that the three-dimensional neuronal chamber is directly underneath and proximal to the epithelial monolayer. The membrane pore size (1 µm) is large enough to support neurite extension through the gel layer and pores. This provides the potential of direct contact between the neurons and epithelium, facilitating both paracrine and synaptic communication. Orthogonal projections of a z-stack collected from our MPS show a distinct neuronal layer (red, beta III tubulin) with some possible extensions through the membrane up to the top epithelial layer expressing ZO1 in green (**Fig. 3e**). A 3D surface plot rendering of the ZO1 and beta III tubulin shows the intensities throughout the z stack of one of our co-cultured MPSs.

### Enteric neuron cultures exhibit a heterogeneous population with cholinergic and VIPergic subtypes potentially modulating the epithelial function

Enteric neurons interact closely with the gut epithelium but are traditionally absent in in vitro gut models. Much like the central nervous system, enteric neurons are composed of several subtypes responsible for different functions within the enteric nervous system. Some of these functions include the production of substances that modulate epithelial barrier formation and differentiation. We investigated the cholinergic and VIPergic neurons in our cultures, which have counteracting effects on the epithelium. Acetylcholine increases the permeability^25^ and decreases the proliferation^26^ of the epithelium. Meanwhile, VIP decreases permeability^23^. Immunocytochemistry was utilized to assess the percentages of ChAT and VIP neurons present in our primary cultures (**Fig. 4a**), which were then quantified in CellProfiler using beta III tubulin staining as a positive control for the neuron boundaries^31^. Our results showed a heterogeneous population of neurons with an average of 31.1% expressing ChAT and 33.4% expressing VIP (**Fig. 4b**). This was broken down further to quantify the ratios of ChAT only and VIP only expressing neurons, with 13.19% expressing VIP only, 10.84% expressing ChAT only, 20.26% expressing both markers, and 55.71% expressing neither (**Fig 4c&d**). Our findings differ slightly from prior literature estimating that the human small intestine contains 5-15% VIPergic and 50-70% cholinergic neurons^34^. Differences could be due to the source organism and maturity, as the experiments presented in this work utilize neonatal rat tissue. Further, many innervating populations reside outside the myenteric plexus. The diversity of enteric neuron subtypes remains understudied. Because we identified our neuron cultures containing a majority VIPergic population, the neurons may contribute to a stronger barrier in the epithelial cells within our co-cultured system.

**Figure 4.**
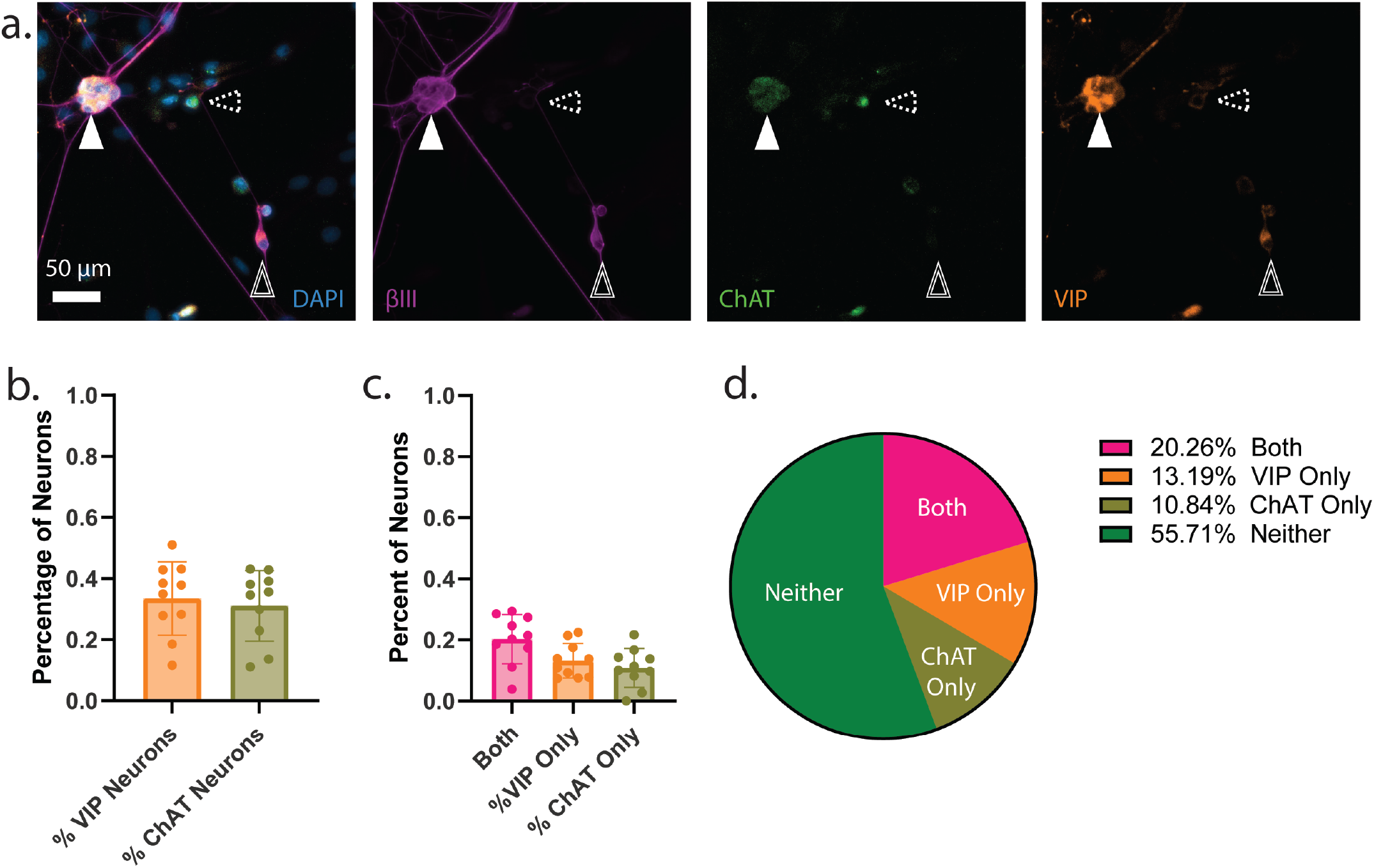
Characterization of myenteric neuron populations utilized on-chip. **a)** Immunocytochemistry images of rat neonatal enteric neurons after 7 days under standard culture conditions. Cultures contained a heterogenous population of VIPergic (VIP, double outlined arrow), cholinergic (ChAT, dashed arrow), and co-expressing neurons (solid white arrow). **b)** About 30% of all the neurons imaged expressed VIP and ChAT. **c, d)** Many neurons expressed both VIP and ChAT in the cultures. Technical Replicates=2 (pooled tissue samples between 10 animals, mixed sex), wells per replicate=4-6.

### Co-Cultured epithelial cells and enteric neurons had more robust barrier properties compared to monocultured epithelium

Our 12-unit MPS system was designed to support real-time on-chip assays for assessing live epithelial barrier properties, including TEER and apparent permeability. TEER was measured using an EVOM device with chopstick electrodes on day 10 (**Fig. 2a**). The design of the medium inlets was made large enough to support easily inserting the electrodes into the chip, similar to a standard Transwell® system (**Fig. 5a**). On a single 12-unit chip, TEER values for the respective monoculture controls, epithelial and ENS co-cultures, and blank controls were measured and compared (**Fig 5b**). As expected, the enteric neuron only group had significantly lower TEER values with an average of 1897.17 Ω/cm^2^ compared to the co-cultured and epithelial cell-only groups at 2345.80 Ω/cm^2^ and 2336.8 Ω/cm^2^ (p=0.0030 and p= 0.0037, respectively). Enteric neuron-only cultures exhibited lower resistances than the blank control (2044.79 Ω/cm^2^), likely because the neuronal cells degraded the gel. Fold change was calculated for each group, comparing the difference between blank and epithelial-only TEER values and enteric neuron-only and co-cultured TEER values to assess the absolute permeability of the epithelial monolayers and control for the effects of the gel degradation. There was a significantly higher fold change in the co-cultured group at 1.25 average fold, compared to the epithelium only at 1.14-fold (p=0.049, **Fig. 5c**).

**Figure 5.**
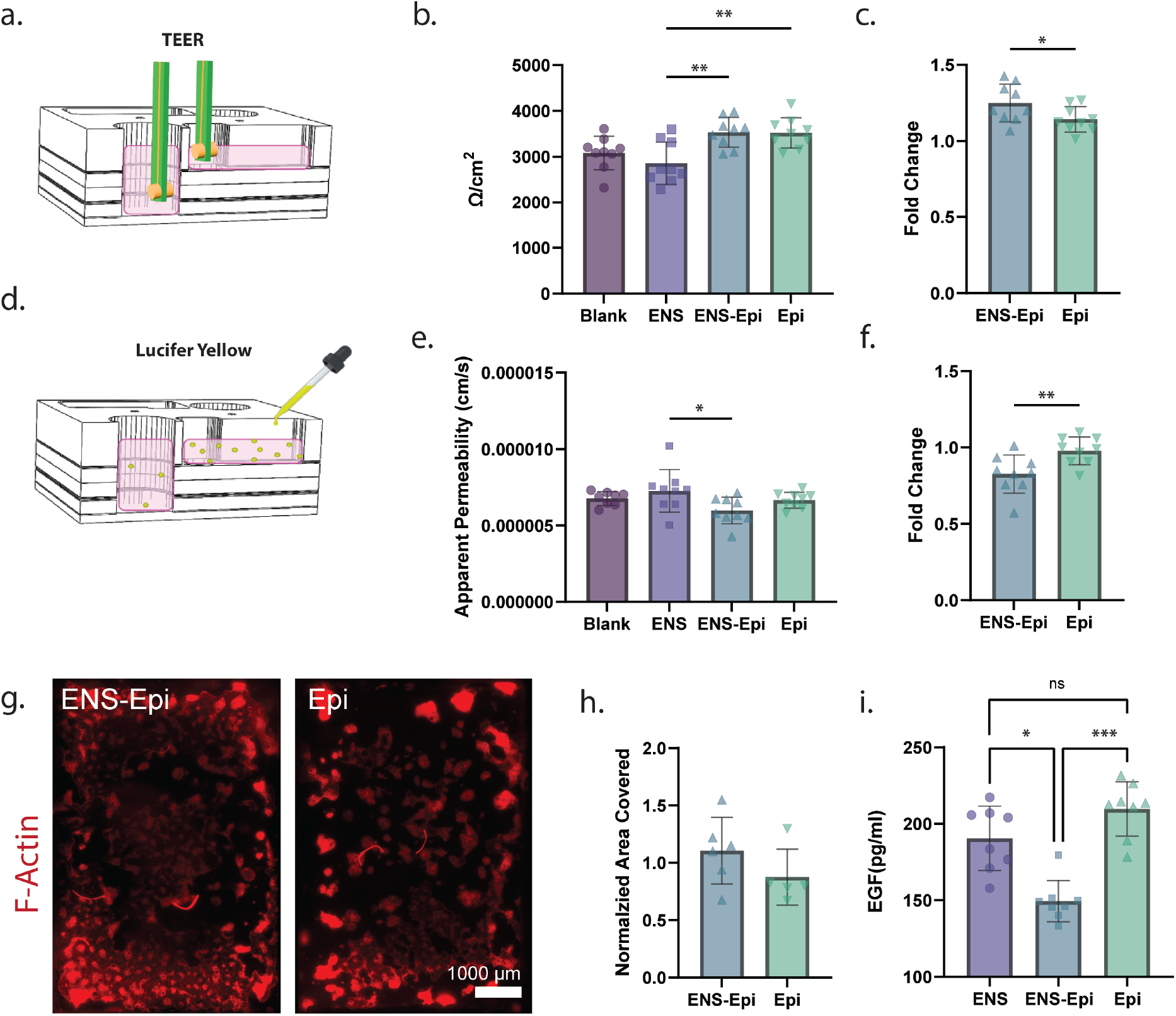
Our MPS system enables traditional barrier strength measures like transepithelial electrical resistance (TEER) and apparent permeability assay. **a)** TEER was measured via chopstick electrodes and the EVOM2. **b)** TEER values were overall greater for cultures containing the epithelium (Epi). **c)** Fold change of the TEER values controlled for the 3D gel layer degrading when ENS was cultured in it. (n=3, m=3) **d)** Apparent permeability, measured by lucifer yellow diffusion through the culture, followed the TEER results. **e)** Less fluorophore traveled through the layers when there was an epithelium present, and **f)** the fold change comparison to appropriate controls showed a significant difference between the epithelium with and without the ENS. (n=3, m=3) **g)** Cell area coverage was also measured using phalloidin-stained monolayers and images of the entire culture area. **h)** The co-culture condition had a trending higher normalized coverage within the culture area than the epithelium alone. (n=2, m=2-3, normalized to an average of all samples in a replicate n). **i)** A higher EGF basal supernatant concentration was recorded from the co-culture condition than either of the monoculture conditions (n=3, m=2-3).

Fluorescent molecule diffusion across the epithelium was measured in parallel to TEER at day 10. Here, we used the small fluorescent molecule lucifer yellow to measure apparent permeability from the apical and basal compartments in our MPS over three hours (**Fig. 5d**). The apparent permeability calculated from lucifer yellow diffusion was significantly lower for the co-culture group (3.97x10-6 cm/s, p=0.042) compared to epithelial cells only (4.41x10-6 cm/s), matching our findings from TEER measurements (Fig. 5e). Fold change was calculated comparing the two control groups (blank and neuron only) to the epithelial cells only and co-cultured groups, respectively. There was a significantly lower permeability fold change of 0.83 for the epithelial and enteric neuron co-culture compared to the epithelial-only monolayer of 0.98-fold (p=0.0093, **Fig. 5f**). Considering that ENS culture degraded the gel, which should lower the barrier to diffusion through the bulk, the ENS improved the epithelial barrier compared to the epithelium alone.

In addition to permeability assays, we quantified the entire area of the MPS epithelial layer coverage by characterizing phalloidin-stained epithelial cells at the same timepoint as barrier function, after 10 days (**Fig. 5g**). The edges of the tiled region are brighter due to reflection of the light from the acrylic. The area covered by cells for each sample was normalized by the average of all sample areas, combining mono- and co-culture, of a full technical replicate (n) to remove some interexperimental variability. The epithelial-only and co-cultured groups were compared, showing a higher normalized area coverage of 1.1048 for the co-cultured group with enteric neurons present compared to epithelial cells only with a mean normalized coverage of 0.8742 (**Fig 5h**, unpaired t test p=0.246). This supports our TEER and Lucifer yellow permeability findings and the hypothesis that enteric neurons promote an improved epithelium barrier function in our in vitro system.

### Co-cultures of enteric neurons and epithelium contain significantly less EGF than their monoculture counterparts

An ELISA assay was run to measure the overall EGF concentration found in mono- and co-culture media supernatants due to the known impact on epithelial barrier function^35,36^. The epithelial media contains 50 ng/ mL of EGF added that either degrades or is consumed over the course of the experiment. For this reason, the blank was not included in the analysis. The co-cultured epithelium and ENS contained significantly less EGF than either monoculture condition (**Fig 5i**). The mean EGF concentration of the co-cultured sample was 149.3 pg/mL, compared to 190.3 and 209.6 pg/mL for neuron only and epithelium only, respectively (p=0.04 and p=0.0003,). This profound difference in EGF in the basal supernatant emphasizes the importance and great differences present in heterogeneous populations in in vitro models.

## Discussion

Understanding the importance of innervation throughout visceral organ systems in the body is an important and understudied field. Most gut-on-chip models or in vitro culture systems lack an enteric neuron component. Here, we developed a gut MPS incorporating a primary epithelial and enteric neuron co-culture. Further, we scaled the platform to 12-units to allow parallelization of our culture conditions to reduce variability via biological factors, human handling, and culture environment. The addition of neurons is critical for developing more complex and biomimetic organ chip devices, especially as they become the standard for studying developmental biology, drug delivery, and disease progression^37^. We have demonstrated key changes in barrier function and paracrine signaling in our system, highlighting the importance of incorporating the autonomic nervous system. The enteric nervous system is heavily involved in immune interactions and is implicated in several neurological and gastrointestinal disorders^38,39^.

In our pursuit, we have overcome several of the current challenges in the robustness, reproducibility, and reliability of organ-chip platforms and developed a device that supports twelve separate co-culture systems for more efficient and higher-throughput experimentation as well as removing the need for tubing and pumps. These MPSs are cost-effective and can be manufactured easily using standard maker space equipment—an improved process from the standard poly(dimethylsiloxane) (PDMS) model and casting methods. Traditional MPSs fabricated via PDMS soft lithography require expensive equipment and expertise in microfabrication, hindering the design/re-design phase and the average lab from adopting the method^40^. Additionally, the process of prototyping multiple models to optimize a MPS design, requiring a few iterations, can be cost prohibitive at $150-500 per design^8^. Our high-throughput, laser cut and assemble chip cost less than $2 per independent culture, can be made in about 3hrs, and can be modified with ease on any computer aided design software. Our fabrication method is flexible, efficient, and accessible to most engineering universities. And our 12 culture MPS can fit four parallel conditions in triplicate on a single device.

The gravity-driven flow was incorporated into the device to make the model more biologically relevant than static cultures, as epithelial cells undergo a lot of shearing in the process of digestion^41,42^. Although in vivo the shearing force is unidirectional. The MPSs support on-chip assays of epithelial barrier function comparable to methods performed in standard Transwell® systems, including fluorescent molecule diffusion and TEER. Although, measures of transport on the MPS cannot be directly compared to Transwell® systems due to the additional membrane layer and 3D gel. Despite the complexity, the system can be imaged on an inverted light microscope, making these an accessible experimentation platform for smaller research operations with limited access to confocal microscopy.

Paracrine and potentially synaptic signaling from the abutting enteric-derived neurons, resulted in a more substantial epithelial barrier as measured with TEER and apparent permeability assays. We speculate that part of this could be due to enteric neurons increasing cell proliferation, barrier integrity, or preventing apoptosis of the epithelium. Enteric neuron subtypes have different primary functions within the gut, and some dominating neuron subtypes may be the reason for decreased epithelial permeability. Our quantification of VIP and ChAT-expressing neurons show a slightly higher percentage of VIP+ cells. These findings, along with the lower permeability seen in our co-cultures, we hypothesize VIP maybe having a greater influence on the culture over acetylcholine. Future work includes dosing in neurotransmitters of interest, like VIP and acetylcholine, to help determine the molecules the neurons might produce and their impact on the epithelial barrier. The enteric neurons may also change the composition of differentiated epithelial cell subtypes, and, as a result, improve the barrier strength. Inclusion of neuron populations from outside the myenteric plexus (i.e., ganglia, spinal cord, and hindbrain) along with further characterization of the epithelial subtypes present in the monolayers could improve our understanding of the direct impact of neuron subtypes on epithelial differentiation.

Previous studies have reported that there is an increase in EGF mRNA when neurons and glia are cultured with the intestinal epithelium^43^. Glia, a support cell type of neurons, are known to produce a precursor to EGF that is less potent but can be activated into EGF to promote healing of barrier tissues when inflamed^35,44^. The epithelium secretes EGF from enterocytes45 and Paneth cells^46^, while the proliferative crypt cells have receptors for the EGF. The increased EGF in the monocultures could be due to the glia and enterocytes producing more in these monocultures, and less (or no) intestinal stem cells consuming the surplus^47^. Alternatively, the lower EGF in co-culture could be due to an increased consumption of the EGF by the intestinal crypt cells. Further characterization of the proliferation of our epithelium and a glia only culture group would help us break down the source and consumption of EGF in our cultures. For the time being, the major takeaway is the enteric nervous system influences the intestinal epithelium and GI models lacking this innervation are missing a vital component to gut renewal.

Although innovative in several areas, including the report of the first GI MPS containing epithelial and neuronal cells, our model has a few limitations. Cells in this model were sourced from neonatal rat cells, which are less representative than a humanized model and potentially less indicative of mature adult tissue^48^. The time frame of these experiments is relatively short compared to what can be achieved with adult human cells, especially in distal intestinal regions. Modeling the duodenum was chosen since it is a heavily innervated part of the intestine and where the bulk of drug absorption occurs,^49^ but it is underrepresented in current literature. However, duodenal organoids and monolayers are not as robust or well documented in literature as those from the colon^50-52^. Colonic tissue has a slower turnover rate and higher levels of mucus producing goblet cells. The slower turnover rate of colonic cells translates to longer-lasting monolayer viability compared to duodenal cells that will differentiate quickly and die off. The increased mucus barrier of the colon also provides a more substantial quantifiable barrier. Incorporating enteric neurons may be a valuable method for improving the quality of in vitro duodenal cultures, expanding the ability to study small intestine disorders.

With the rapid discovery new connections between the brain and the gut daily, creating models of the neurological links to the gut in a reductionist manner is paramount in finding the underlying mechanisms. Our results display that enteric neurons influence the function and health of the intestinal epithelium without an exogenous or inflammatory challenge. This work proves that future steps should strive to include more complex tissues like ENS. The model established and characterized in this work is the first innervated small intestinal MPS. This work is a jumping-off point for future studies of neuronal communication to and from the gut.

## Methods

### Chip design and fabrication

Innervated MPSs were assembled layer-by-layer using laser cut thermoplastics with our established method, however we scaled the system to include twelve individual “chips” onto one device^8,27^. Transparent, cast polymethyl methacrylate (PMMA, McMaster-Carr, 8560K211 and 8560K171) and clear 0.005-inch polyethylene terephthalate (PET) film (McMaster-Carr, 8567K54) were used for the cell chamber wall layers. Tissue culture-treated PET, semi-permeable membranes with 1 um pores (*It4Ip, ipCellCulture 2000M12/620M103*) were utilized to separate the cell populations. Layer assembly was done using pressure-sensitive transfer tape (3M, 966), and the chip was built on top of a glass slide base layer (Ted Pella, 260230-50). Designs for each MPS layer were created as vectors in Adobe Illustrator and cut using an Epilog Zing laser cutter. Transfer tape was laminated onto the desired layers before laser cutting. MPSs were assembled by hand, heat pressed and then placed into a vacuum oven for at least 120 hours at 50°C to allow curing and off-gassing of the adhesive material. Binder clips were used to apply additional pressure to the newly assembled MPS while baking.

### Animal Care and Use

The following tissue isolation procedures were approved by Northeastern University’s Institutional Animal Care and Use Committee (IACUC). All methods were preformed following the guidelines and regulations necessary.

### Primary Neuron Isolation

Enteric neurons were isolated from neonatal two-day-old rats (p2) sacrificed via decapitation. The small intestines are removed, and the myenteric plexus was carefully peeled off the intestines using forceps under a dissecting microscope and stored in ice-cold Hanks Balanced Salt Solution (HBSS). The peeled myenteric plexus was moved into a 15 mL conical tube and digested for one hour in collagenase type II (Gibco, 17101015, 1 mg/mL) and Deoxyribonuclease I (Sigma, 0.5 mg/ ml) dissolved in neurobasal medium (Gibco, 10888022) at 37°C. The myenteric plexus in the digestion medium was vortexed for 5 seconds and examined visually. The digestion continued in 30-minute increments until the myenteric plexus was visually broken into small particles. Usually, this is complete after three additional 30-minute digestion steps. The solution was then centrifuged at 500 g for 5 minutes, and the supernatant was removed. The pellet was resuspended in a 0.05% trypsin-EDTA (Gibco, 25200056) solution for 30 minutes at 37° C to dissociate the neurons. After dissociation, the solution was centrifuged at 500 g for 5 minutes, and the trypsin supernatant was removed. The cell pellet can be resuspended in enteric neuron medium (**Table 1**) and counted for culture. The fully digested myenteric plexus should only have neurons remaining and will look like a rectangular grid structure under the microscope. The fully digested neural tissue can be seeded directly onto treated glass or well plates and fed with enteric neuron medium (**Table 1**) or moved into nonadherent well plates for the culture of neurospheres. Neurospheres can then be kept in suspension culture with enteric neuron media addition without any growth factors added every two to three days.

**Table 1:** Enteric neuron medium composition.

| Medium Reagent | Supplier/Catalog # | Final Concentration |
| --- | --- | --- |
| Neurobasal Medium | Neurobasal-A, 10888022 | 97% |
| Fetal Bovine Serum | Corning, 35015CV | 1% |
| Antibiotic/Antimycotic | Gibco, 15240-062 | 1% |
| Glutamax\| | Gibco, 35050061 | 1X |
| GDNF | Gibco, PHC7045 | 10ng/mL |
| NGF | R&D Systems, 256-GF/CF | 25ng/mL |

### Epithelial Organoid Isolation

Duodenal epithelial organoids were isolated from the first 3 - 4 cm of the small intestine of neonatal rat pups (p2). After the myenteric plexus was removed from the segment, the remaining intestine was fileted open and cut up into small pieces (∼5 mm). The diced intestines were washed ten times with cold phosphate-buffered saline (PBS), and with the final wash, the PBS was removed and replaced with 2 mM EDTA (Invitrogen, AM9260) in PBS. The tissue was left to digest on ice and rocking for 15 minutes. The tissue was then allowed to settle, and the EDTA solution was removed and replaced with fresh 2 mM EDTA and digested for an additional 25 minutes on ice with rocking. To detach the crypts from the rest of the tissue, the tube was vortexed for 1 minute, followed by pipetting with a 10 mL serological pipette while mixing about 25 times. The suspension was filtered through a 100 µm cell strainer and spun down at 300g for 5 minutes. The supernatant was removed, and the pelleted crypts were resuspended in Advanced DMEM/F12 (Gibco, 12634028) and spun again to wash, this was repeated three times. After the final wash and suspension removal, 20 µL of crypts were encapsulated in 1 mL growth factor reduced Matrigel® (Corning, 356231) and then plated into 50 µL domes in a 12-well plate, three domes per well. After the Matrigel® had polymerized, the cells were fed an expansion medium (**Table 2**) with an addition of 10 µM rock inhibitor, Y-27632. Organoids were fed every 2-3 days with fresh expansion medium and passaged every 5-7 days.

**Table 2:** Expansion medium composition.

| Medium Reagent | Supplier & Catalog # | Working Concentration | Final Concentration | Volume for 50 mL |
| --- | --- | --- | --- | --- |
| WRN Media | HDDC Organoid Core/Breault Lab | N/A | 50% | 25 mL |
| Adv DMEM/F12 | Gibco, 12634028 | N/A | 45% | 22.5 mL |
| Glutamax | Gibco, 35050061 | 100X | 1X | 500 µL |
| HEPES | Fisher, AAJ16924AE | 1 M | 10 mM | 500 µL |
| Primocin | Invivogen, NC9141851 | 50 mg/mL | 100 µg/mL | 100 µL |
| Normocin | Invivogen, NC9273499 | 50 mg/mL | 100 µg/mL | 100 µL |
| B27 | Gibco, 17504-044 | 50X | 0.5X | 500 µL |
| N2 | Gibco, A13707-01 | 100X | 0.5X | 250 µL |
| N-Acetyl-Cysteine | Sigma, A7250 | 500 mM | 500 µM | 50 µL |
| A-83-01 | Sigma, SML0788 | 500 µM | 500 nM | 50 µL |
| SB202190 | Sigma, S7067 | 30 mM | 10 µM | 16.6 µL |
| EGF | Peprtech, 315-09 | 500 µg/mL | 50 ng/mL | 5 µL |
| Gastrin | Sigma, G9145 | 100 µM | 10 nM | 5 µL |
| Y-27632 (only added after passaging) | Tocris\|Bioscience, 12-541-0 | 10 mM | 10 µM | 50 µL |

Organoids were passaged by removing the media, replacing it with Cell Recovery Solution (Corning, 354270), and then disrupting the Matrigel® domes by scraping them off the plate with a pipette tip. Organoids, Matrigel® fragments, and cell recovery solution were collected into a tube and left on ice rocking for 45 minutes to degrade the Matrigel®. Organoids were then spun down at 500 g for 5 minutes, and the supernatant was removed. At this point, 20 µL of the pellet was collected and resuspended in 1 mL of Matrigel®, or the cell underwent further digestion for monolayer seeding.

### MPS Seeding

Fabricated co-culture chips were sterilized by UV for ten minutes, followed by oxygen plasma treatment for 90 seconds. The top membrane was then coated with poly-d-lysine (Gibco, A3890401) and incubated for 1 hour at 37° C, and then rinsed with PBS at least twice. Next, a 1:10 dilution of Matrigel® coating was added at 150 µL in preparation for epithelial monolayer adhesion and incubated for at least 1 hour at 37° C before seeding chips with the three-dimensional enteric neuron culture. To seed organ chips with the neurospheres, the enteric neuron media with suspended neurospheres was first collected from the wells using a 1000 µL pipette and placed into a 15 mL tube. The tube was centrifuged at 300g for five minutes to pellet the neurospheres. The supernatant was removed, and 1 mL of Accutase (Corning, 25058CI) was added to the pellet. The Accutase solution was triturated to break up the pellet, and the tube was incubated for 30 minutes at 37° C. After incubation, the neurospheres were triturated 25-45 times using a 1 mL pipette tip and centrifuged at 300g for 5 min. The Accutase was removed and replaced with fresh neuron medium, and cells were counted and seeded on-chip at 1x106 cells/mL in a 10% Matrigel® (Corning, 356231), 2 mg/mL Cultrex® Collagen (R&D Systems, 344702001) solution in neuron media. The volume of cells needed to achieve 1x10^6^ cells/mL was mixed with the Matrigel® and Cultrex® solution and 10 µL was injected into inlets on the MPS that lead into the enteric neuron chamber. The gels were thermally crosslinked at 37° C for 20 minutes before adding medium to the reservoirs of the chip. Neurons received a complete media exchange after 24 hours with 20 µM of the mitotic inhibitor, cytosine β-D-arabinofuranoside (AraC, Sigma, C1768). During the remaining timeframe, the neurons received half-volume media exchanges.

When ready to seed an epithelial monolayer, the Matrigel® domes were disrupted with manual scraping using a pipette tip. Organoids and Matrigel® fragments were collected in cell recovery solution and rocked on ice for 45 minutes to remove the organoids from the Matrigel®. Organoids were centrifuged at 500g for 5 minutes, and the cell recovery solution was removed. Organoids were resuspended in trypLE (Gibco, 12563) and digested for 5 minutes at 37° C. The cells were then agitated by pipetting up and down through a bent 1 mL pipette tip 20 times to achieve a single-cell suspension. Cells were spun down to remove the trypLE, resuspended in the expansion medium with 10uM Y-27632 rock inhibitor, and counted. Chips were seeded with 600,000 cell/chip, 2.2×10⁶ cells/cm^2^. After one day, the media was changed out for differentiation medium (**Table 3**).

**Table 3:**
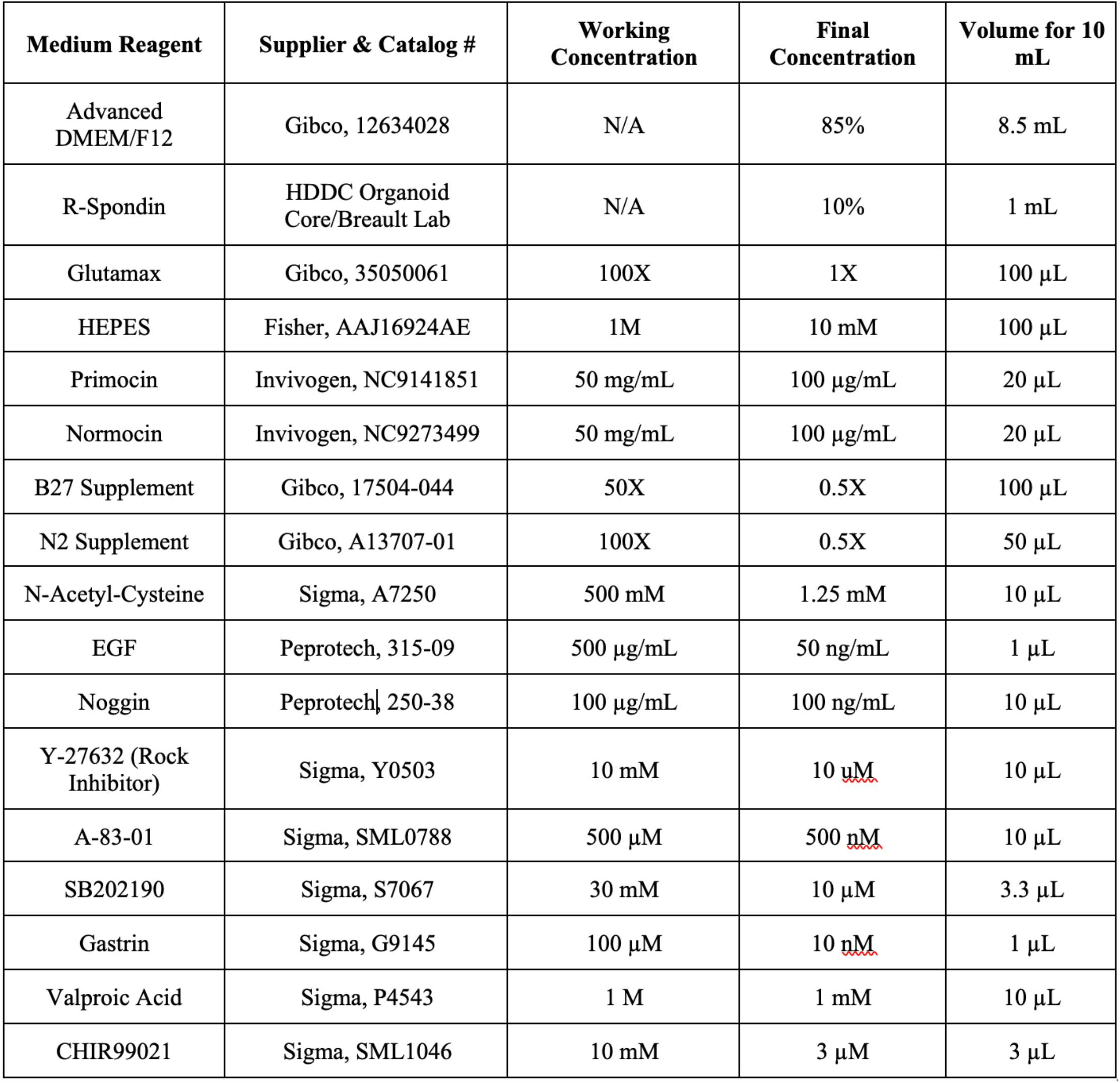
Epithelial differentiation medium composition.

### Media Compositions

According to supplier suggestions, lyophilized reagents were resuspended to their working concentration in deionized water, DMSO, or PBS. Rat enteric neuron medium was used for feeding neuron cultures (**Table 1**), with a volume half exchange performed every two to three days. Rat organoid proliferation medium (**Table 2**) was used during organoid expansion in Matrigel® domes and for the first 24 hours of seeding a monolayer. For chip monolayer experiments, after 24 hours, the epithelial medium reservoir was exchanged for rat organoid differentiation medium (**Table 3**) to encourage differentiation into representative heterogeneous cell populations.

### Application of Shear Stress on-chip

MPSs were placed on a platform rocker (ThermoScientific, 11-676-680) in an incubator with 5% CO2 at 37° C for three days. A rocker tilt angle of 2° and rotation speed of 10 rpm were determined to generate a physiological rate of shear force (0.002–0.08 dyne/cm^2^), calculated at the center of the monolayer culture surface area when at the maximum tilt angle^6^. Calculations were determined using the assumptions and formulas stated in a previous publication^28^. The critical flip angle, *θ*_*0*_, when the fluid-free surface first contacts the edge of the bottom of the MPS, was calculated using the MPS’s epithelial channel dimensions (**Eq. 1**). Here, *h*_*0*_ represents the medium depth, and *L* represents the length of the epithelial channel. Once the critical flip angle was determined, the shear stress could be calculated using Equation 2, where *θ* is the rocking angle, *θ*_*0*_ is the critical flip angle, *T* is the period, and *x* is the location of interest within the channel.

### Permeability of Lucifer Yellow

Lucifer yellow lithium salt (Invitrogen, L453) was resuspended to a 100 µM stock solution in PBS and wrapped in foil to avoid light exposure. A 10 mM working solution was prepared in phenol red free DMEM (Gibco, 31053028). All media was carefully removed from apical and basal chip compartments, and 150 µL of 10 mM Lucifer yellow was added into the apical compartments of all the chip wells. Phenol red free DMEM was added to the basal compartments (150 µL) to avoid drying the neurons out. The chips were incubated at 37°C for three hours at static conditions. After incubation, 100 µL of the solution was removed from the apical and basal channels and put into separate wells of a black 96-well plate. A standard curve for the Lucifer yellow working solution was prepared using 1:2 serial dilutions of the 10 mM solution and phenol red free DMEM. The fluorescence was then measured for each well on a plate reader spectrophotometer using 428/530 nm excitation/emission settings with the gain set between 50-75. Apparent permeability was then calculated using the concentration values determined from a linear fit of the standard curve.

### Immunostaining

Cells were fixed with paraformaldehyde for ten minutes. After fixation, the solution was removed from the culture chambers, and a rinse with HBSS was performed twice. Cells were permeabilized in a 0.1% Tween-20 solution for ten minutes. After rinsing out the Tween, a blocking solution of filtered 4% goat serum (Sigma, G9023) was added to the culture and left to incubate overnight at 4° C. The following day, the blocking solution was removed. Antibodies (**Table 4**) diluted in 4% goat serum were added into the wells and incubated for two hours at room temperature or overnight at 4° C. Following antibody addition, MPS compartments were carefully rinsed with HBSS three times. Secondary antibodies diluted in 4% goat serum were then added and incubated for two hours at room temperature or overnight at 4° C. Following secondary incubation, DAPI (Invitrogen, D1306) was added at 1:1000 dilution in PBS for 10 minutes. Rinsing was then performed carefully in the MPS at least three times. MPSs were imaged immediately or stored at 4° C and wrapped with parafilm.

**Table 4:** Antibodies and dilutions used for immunocytochemistry.

| **Antibody** | **Catalog ID/Supplier** | **Host** | **ICC Dilution** | **Notes** |
| --- | --- | --- | --- | --- |
| Beta III Tubulin | Sigma, T8660 | Mouse | 1:1000 | Structural, mature neurons |
| Zonula Occludens-1 (ZO-1) | Invitrogen, 40-2200 | Rabbit | 1:100-1:200 | Tight junctions, mature epithelium |
| Vasoactive Intestinal Peptide (VIP) | Abcam, AB8556 | Rabbit | 1:200 | VIP producing neurons |
| Choline Acetyltransferase (ChAT) | Millipore, AB15468 | Chicken | 1:1000 | Acetylcholine producing neurons |
| Phalloidin Alexa Fluor 647 | Invitrogen, A22287 | NA | 1:1000 | F-actin |
| Anti-Chicken, Alexa Fluor 488 | Invitrogen, A11039 | Goat | 1:1000 | Secondary |
| Anti-Rabbit, Alexa Fluor 546 | Invitrogen, A11035 | Goat | 1:1000 | Secondary |
| Anti-Mouse, Alexa Fluor 647 | Invitrogen, A32728 | Goat | 1:1000 | Secondary |

### Terminal Epithelial Monolayer Characterization

Monolayers were stained for filamentous actin with Phalloidin to identify the cell cytoskeleton and total area. Tiled images of the entire culture area of the immunostained MPSs were acquired on a Zeiss Axio Observer Z1 (Carl Zeiss Microscopy). Exported images were analyzed using Fiji in ImageJ^29^. Briefly, the images were cropped to a uniform size matching the dimensions of the chip culture area. Then a variance filter was applied to improve contrast at the edges of the cells. The MorphoLibJ plugin was used to segment the image, and the total cell area covered was recorded^30^.

### Acetylcholine and Vasoactive Intestinal Peptide Neuron Subtype Quantification

Neonatal rat enteric neurons were isolated as described above and plated on cover glasses. Cover glasses were sterilized by UV for 5 minutes on each side, oxygen plasma treated for 90 seconds, poly-d-lysine (Gibco, A3890401) coated for 1 hour followed by washes, and finally 50 µg/mL mouse laminin (Corning, 354232) coating for 30 minutes. Neurons were seeded at a density of 50,000 cells/well in a 24-well plate with a half-exchange medium feeding every two to three days.

After 1 week, the neurons were fixed with 4% paraformaldehyde for ten minutes and then washed with HBBS 3 times. The cells were permeabilized with 0.01% Triton-X for ten minutes, followed by additional washes with HBBS. Blocking was done with 2.5% goat serum (Sigma, G9023) for at least 1 hour before adding the primary antibodies for Beta III Tubulin, VIP, and ChAT (see product numbers and dilutions in Table 4). Primary antibodies were incubated at room temperature for at least an hour, followed by 3 washes with HBBS. The secondary antibody solution was similarly made in 2.5% goat serum and incubated at room temperature for at least an hour, followed by 3 washes. Cover glasses were then mounted on microscope slides with ProLong Gold Antifade Mountant and DNA Stain DAPI (Invitrogen, P36931).

Images of the stained neurons were segmented and quantified using Cell Profiler^31^. Regions of interest were segmented using the nuclei stain of DAPI as a seed and Beta III Tubulin for the boundaries of the cell. Individual cells were quantified for their integrated intensity, the sum of pixel intensities per region of interest. They were considered positively stained for VIP or ChAT if they were greater than 1 standard deviation from the mean of all the neurons imaged per experimental replicate. Immunostained samples were imaged on a Zeiss Axio Observer Z1 (Carl Zeiss Microscopy) for the MPS z-stacks, MPS orthogonal projections, neuronal subtype and phalloidin cell coverage experiments.

### Transepithelial Electrical Resistance (TEER) Measurements

TEER was performed using a World Precision Instruments EVOM device with chopstick electrodes. Electrodes were first cleaned in 70% ethanol and calibrated in PBS. Recordings were made by completely submerging the electrodes into the media of the apical and basal chambers and allowing the resistance reading to equilibrate. A recording was taken for each basal inlet of each sample. The final TEER values for each sample were the average of the two recordings. TEER calculations were then normalized to the area of the chip (0.49 cm2) to get Ω/cm^2^.

### EGF Secretion

At experiment end points, cell culture supernatants were collected separately from each sample’s apical and basal compartments and centrifuged at 3000 rpm for 10 minutes at 4°C. The centrifuged supernatant was then moved into a clean microcentrifuge tube and stored immediately at -80°C until needed. An EGF ELISA (Invitrogen, EREGF) was performed on the collected basal supernatant following supplier instructions with samples ran in duplicate and averaged. Optical densities were obtained using a plate reader at 620 nm and compared to a standard curve fit.

### Statistical Analyses

GraphPad Prism was used to run statistical testing. Normality tests were performed on each data set. Data sets were then analyzed for statistical significance (p<0.05) using an upaired t-test or one-way ANOVA with multiple comparisons for normally distributed data sets and a Kruskal-Wallis nonparametric test for non-normal data sets. All experiments, excluding the phalloidin area coverage, were performed independently in triplicate. Litters of 10 rat pups were used and pooled per experimental trial.

## Acknowledgements

The authors would like to thank Dr. Adam Bindas for the preliminary experimental work that fed into the conception of this work. Thank you to Bryan Schellberg for assistance in designing the gravity-driven flow chip. Figure schematics were partly created using BioRender.com.

## Author contributions

K.N. and J.S. contributed equally to planning, experiments, and manuscript writing. A.K., K.N., and J.S. conceived the experiments. K.N. and J.S. ran the experiments, analyzed the data, and composed the manuscript. A.K. and R.K. provided feedback on experimental design, analysis, and manuscript preparation.

## Supplemental Data Tables

